# Unveiling Novel Double-Negative Prostate Cancer Subtypes Through Single-Cell RNA Sequencing Analysis

**DOI:** 10.1101/2023.08.11.553009

**Authors:** Siyuan Cheng, Lin Li, Yunshin Yeh, Yingli Shi, Omar Franco, Eva Corey, Xiuping Yu

**Author notes:** Correspondence to S. Cheng and X. Yu.

## Abstract

Recent advancements in single-cell RNA sequencing (scRNAseq) have facilitated the discovery of previously unrecognized subtypes within prostate cancer (PCa), offering new insights into disease heterogeneity and progression. In this study, we integrated scRNAseq data from multiple studies, comprising both publicly available cohorts and data generated by our research team, and established the HuPSA (<u>H</u>uman <u>P</u>rostate <u>S</u>ingle cell <u>A</u>tlas) and the MoPSA (<u>M</u>ouse <u>P</u>rostate <u>S</u>ingle cell <u>A</u>tlas) datasets. Through comprehensive analysis, we identified two novel double-negative PCa populations: KRT7 cells characterized by elevated KRT7 expression, and progenitor-like cells marked by SOX2 and FOXA2 expression, distinct from NEPCa, and displaying stem/progenitor features. Furthermore, HuPSA-based deconvolution allowed for the re-classification of human PCa specimens, validating the presence of these novel subtypes. Leveraging these findings, we developed a user-friendly web application, “HuPSA-MoPSA” (https://pcatools.shinyapps.io/HuPSA-MoPSA/), for visualizing gene expression across all newly-established datasets. Our study provides comprehensive tools for PCa research and uncovers novel cancer subtypes that can inform clinical diagnosis and treatment strategies.

**Graph abstract:** 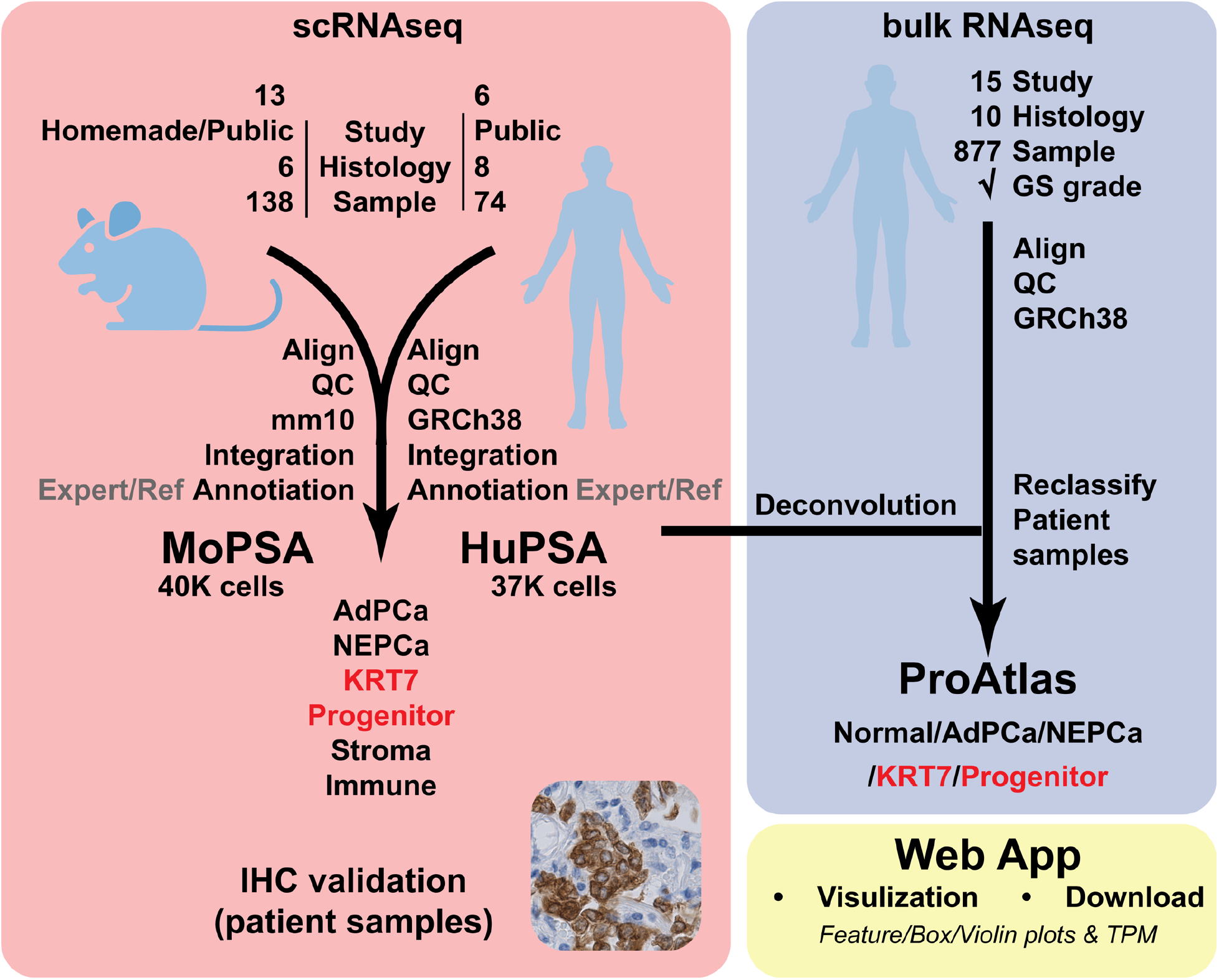

## Introduction

Prostate cancer (PCa) poses a substantial public health burden in the United States. While new AR signaling inhibitors have significantly improved patient survival, these treatments eventually fail, leading to the development of castration-resistant PCa (CRPCa)^1^. About 50% of CRPCa cases retain the characteristics of prostate adenocarcinoma (AdPCa), expressing markers associated with the luminal epithelia of the prostate, such as androgen receptor (AR), AR targets, and HOXB13^2,3^. Approximately 30% of CRPCa present an alternative phenotype known as neuroendocrine prostate cancer (NEPCa). NEPCa cells express minimal levels of AR and its downstream genes, while expressing high levels of NEPCa markers such as CHGA and NCAM1^4^. In addition to the aforementioned phenotypes, there is a less studied subtype known as double-negative prostate cancer (DNPCa), which is characterized by the absence of AR signaling and NEPCa features^5,6^.

Lineage plasticity is implicated in the transition from AdPCa to NEPCa and DNPCa. This phenomenon empowers tumor cells to undergo alterations in their morphology and functions to adapt to challenging environmental conditions, allowing them to transit into new cellular lineages^7^. Key molecular players implicated in this process include the reduced expression of the prostate lineage marker HOXB13^3^ and the aberrant expression of several genes, including SOX2, FOXA2, SRRM4, ONECUT2, BRN2 and PEG10^6-12^. Additionally, the activation of JAK/STAT signaling pathway has been identified as a pivotal contributor to lineage plasticity in PCa progression^6-12^.

Single-cell RNA sequencing (scRNAseq) has emerged as a powerful tool for dissecting tumor heterogeneity. Previous studies have identified various cell populations within human and mouse prostatic tissues, including AdPCa, NEPCa, luminal and basal epithelia, stromal, and immune cells^6,13-27^. However, limitations such as sample size and model-specific populations persist.

To address these limitations, we integrated scRNAseq data from multiple studies and established the Human Prostate Single cell Atlas (HuPSA) and the Mouse Prostate Single cell Atlas (MoPSA). Leveraging these comprehensive datasets, we aimed to identify and characterize known and novel cell populations within prostatic tissues, shedding light on PCa heterogeneity and progression.

## Results

### 1. Establishment of HuPSA and MoPSA

Using scRNAseq data derived from multiple studies, we constructed two combined datasets the Human Prostate Single cell Atlas (HuPSA) and the Mouse Prostate Single cell Atlas (MoPSA). The HuPSA dataset compiled single cell RNA sequencing data of 74 distinct samples from 6 studies, consisting of 368,831 high-quality cells. These samples represent eight histology groups, including normal, normal adjacent, benign, AdPCa, CSPCa (castration sensitive PCa), CRPCa, mCRPCa (metastatic CRPCa), and Cribriform PCa. For the establishment of MoPSA, we generated scRNAseq data from a TRAMP tumor, a widely utilized mouse model for NEPCa research, and combined it with publicly available mouse scRNAseq data. The compiled MoPSA dataset consisted of 395,917 high-quality cells originated from 138 samples obtained from 13 studies. These murine prostatic samples were classified into six groups: WT (wild-type), GEM (genetically engineered mice), control (GEM control mice that express tamoxifen-induced YFP in prostate), castration, regeneration (androgen -induced prostate regeneration), and development (developing prostate)

### 2. Analysis and annotation of cell clusters

UMAP dimensional reduction and unsupervised clustering analysis of the scRNAseq data revealed discernible cell clusters, including luminal, basal/basal-like, progenitor-like (in HuPSA only), NEPCa, and non-epithelial clusters (Figs. 1A & 1B). These clusters were further annotated into distinct populations, primarily based on the expression of marker genes, with considerations of sample histology and inferred copy number variation (iCNV) patterns. This results in 26 and 21 populations identified within the HuPSA and MoPSA datasets, respectively (Fig. 1C & sFig. 1). For a detailed description of each cell population, refer to the supplemental information. Sankey diagrams were used to visualize the data flow between these cell populations and the histology of samples (sFig. 1A), highlighting the widespread distribution of different cell types across diverse histological groups and sample sources.

**Fig. 1.**
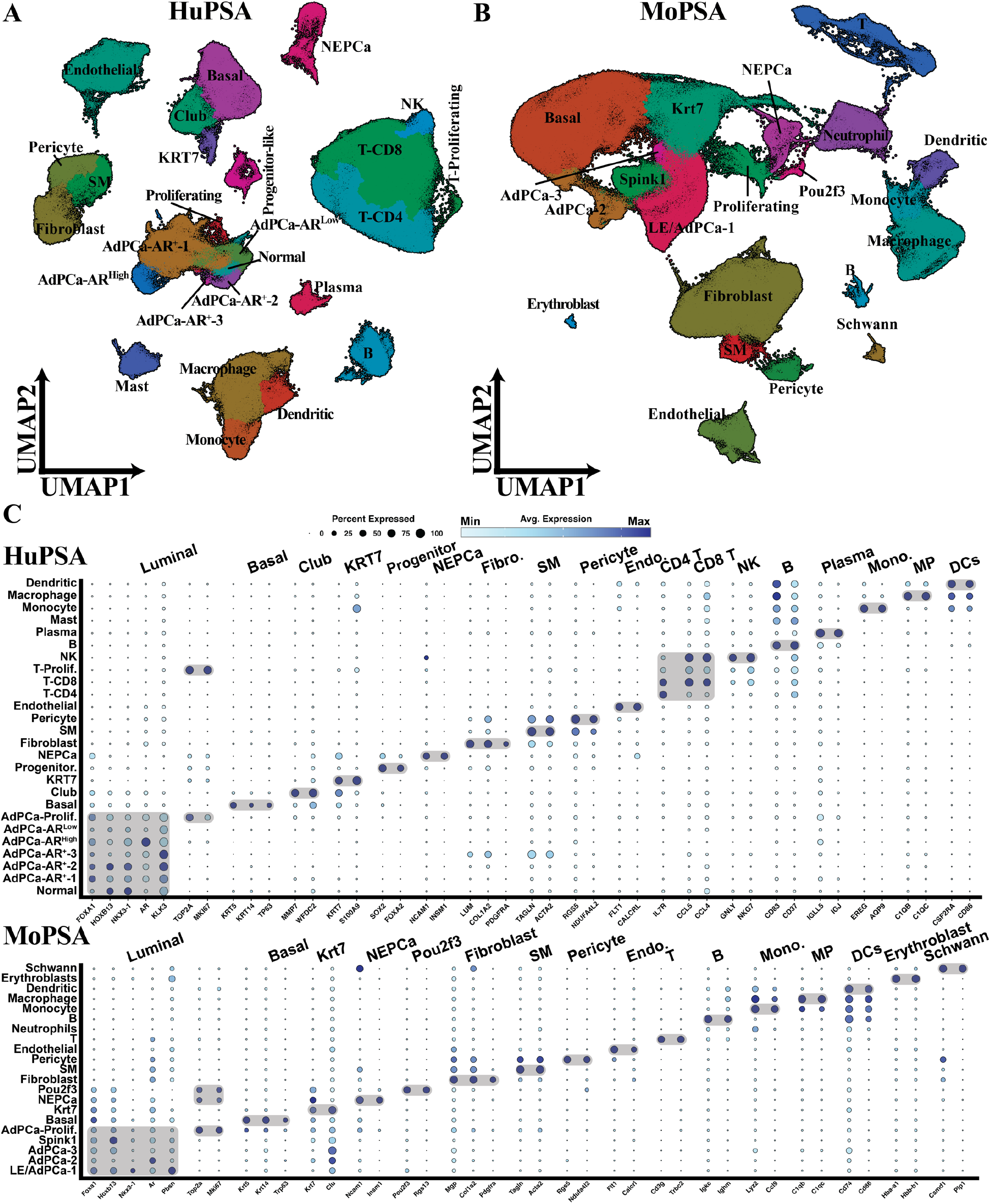
Overview of HuPSA and MoPSA populations. (A) UMAP dimensional reduction plot displaying 26 cell populations in HuPSA. (B) UMAP dimensional reduction plot displaying 21 cell populations in MoPSA. (C) Dot plots illustrating the top marker genes of each cell population identified in HuPSA and MoPSA.

The luminal clusters were characterized by the expression of prostatic luminal epithelial markers FOXA1, HOXB13, and NKX3-1 (Fig. 1C). It was present in both HuPSA and MoPSA, comprising multiple populations that expressed AR at various levels (Figs. 1A, 1B & sFig. 2).

In HuPSA, the basal/basal-like clusters comprised three populations (Figs. 1A, 1C, sFig. 3): basal (KRT5^+^, KRT14^+^, TP63^+^), club (MMP7^+^, WFDC2^+^), and KRT7 (KRT7^+^, S100A9^+^). In MoPSA, the basal and Krt7 populations were observed, while the club group was absent (Fig. 1B).

NEPCa cells from both HuPSA and MoPSA expressed NEPCa markers INSM1 and NCAM1 (Fig. 1C). While mouse NEPCa cells exhibited relative uniformity, human NEPCa cells could be further divided into distinct sub-populations based on the expression of marker genes or enriched pathways (sFig. 4).

In addition to epithelial populations, both HuPSA and MoPSA captured stromal, endothelial, and immune cells as well (Figs. 1A & 1B). Notably, a large population of T cells were detected in HuPSA, predominantly contributed by normal adjacent and AdPCa samples (Fig. 1A & sFig. 1A), consistent with the observation that advanced PCa is an immunologically “cold” tumor.

### 3. Identification of two novel double negative PCa populations: KRT7 and progenitor-like

Our analysis unveiled two novel double negative (DN)PCa populations, namely KRT7 and progenitor-like.

The KRT7 population within HuPSA exhibited heightened expression of KRT7 and several S100 family genes. Despite being clustered together with basal epithelia, this population showed minimal expression of canonical basal markers (Fig. 1C). The KRT7 population comprised 1937 cells, 1828 of them (94%) originating from mCRPCa samples. Additionally, this population displayed elevated levels of iCNVs compared to normal cells or early-stage AdPCa (sFig. 1B). These features strongly suggest that the KRT7 population predominantly comprises cancerous cells rather than nonmalignant basal epithelial cells.

Moreover, the KRT7 population exhibited minimal expression of NEPCa markers and low AR/AR-signaling activity (Fig. 2A), indicating its classification as a DNPCa subtype.

**Fig. 2.**
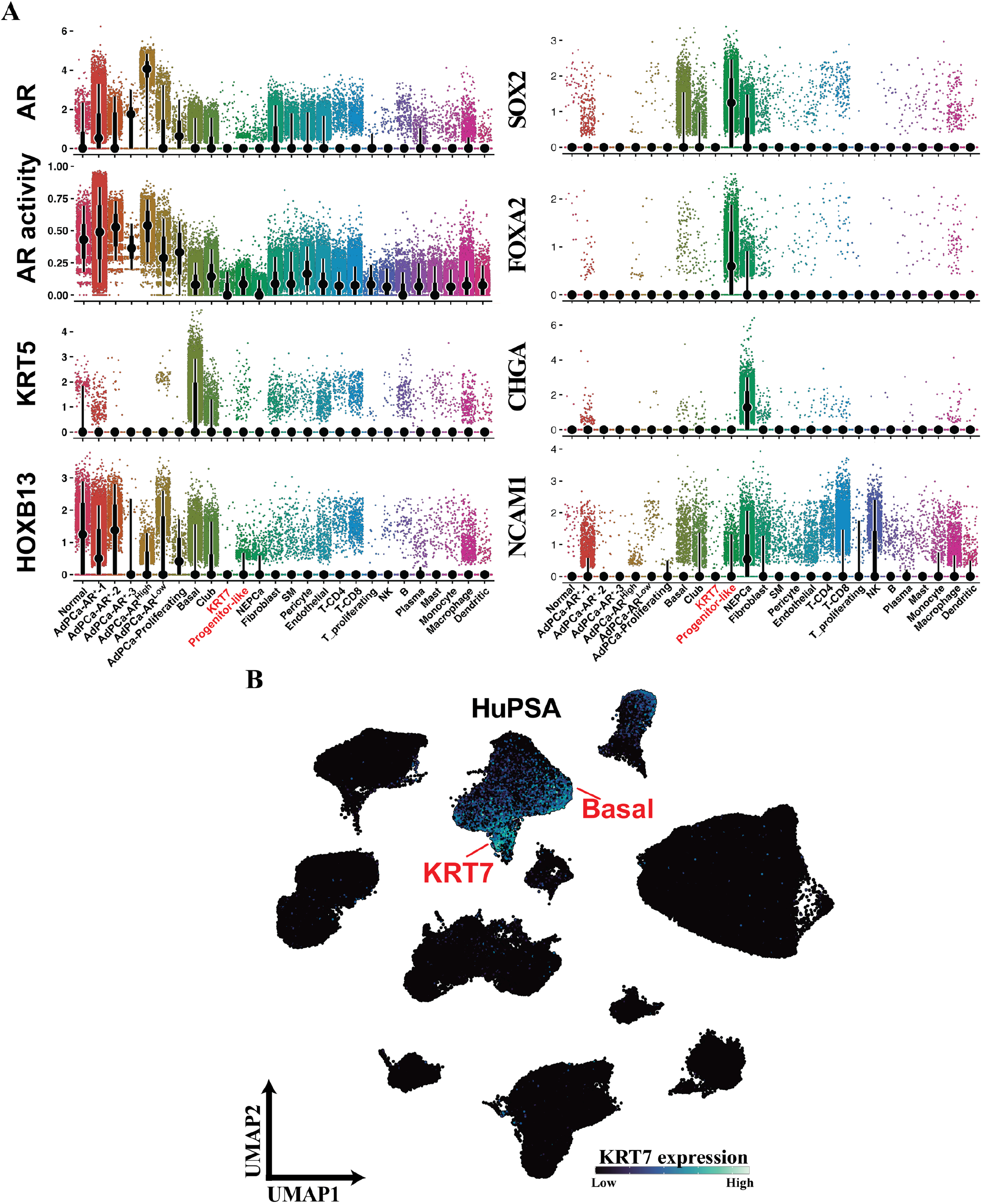
Characterization of KRT7 and progenitor-like populations molecular features. (A) Geyser plots visualizing the expression of AR, KRT5, HOXB13, AR signaling activities, FOXA2 SOX2, CHGA and NCAM1 among different cell populations in HuPSA. KRT7 cell population were AR-negative and basal markers-negative. Progenitor-like cells exhibited expression of FOXA2 and SOX2 but lacked expression of other NEPCa markers CHGA and NCAM1. Moderate expression of SOX2 and FOXA2 was also detected in NEPCa cells. (B) Feature plot illustrating the enrichment of KRT7 expression in both basal and KRT7 populations.

Furthermore, it expressed minimal levels of prostate lineage and differentiation markers, such as FOXA1, HOXB13, and NKX3-1 (Figs. 1C & 2A), indicating a poorly differentiated state.

The progenitor-like population, positioned between the AdPCa and NEPCa populations on UMAP, was exclusively identified in HuPSA (Fig. 1A). Primarily originated from mCRPCa samples, cells within this population exhibited high iCNVs, consisting with their late-stage PCa status (sFig. 1B). Termed “progenitor-like”, this population displayed high expression of SOX2 and FOXA2 (Fig. 2A), typically expressed in embryonic prostate cells rather than in differentiated prostate luminal epithelial or AdPCa cells^28,29^. Notably, NEPCa cells also expressed SOX2 and FOXA2^28,29^ (Fig. 2A). But unlike NEPCa, the progenitor-like cells showed minimal expression of NEPCa marker genes CHGA and NCAM1 (Fig. 2A). Similar to KRT7, progenitor-like cells exhibited minimal expression of AR, AR targets and luminal epithelial lineage genes, supporting a stem/progenitor-like phenotype (Figs. 1C & 2A).

### 4. KRT7 and progenitor-like PCa exist in human specimens

As the discovery of novel KRT7 and progenitor-like DNPCa subtypes emerged from HuPSA analysis, it became imperative to validate their presence in an external collection. Therefore, we compiled publicly available bulk RNAseq data that were derived from 877 human prostatic specimens^30^ and established the ProAtlas (<u>P</u>rostate <u>A</u>tlas) dataset (Fig. 3A).

**Fig. 3.**
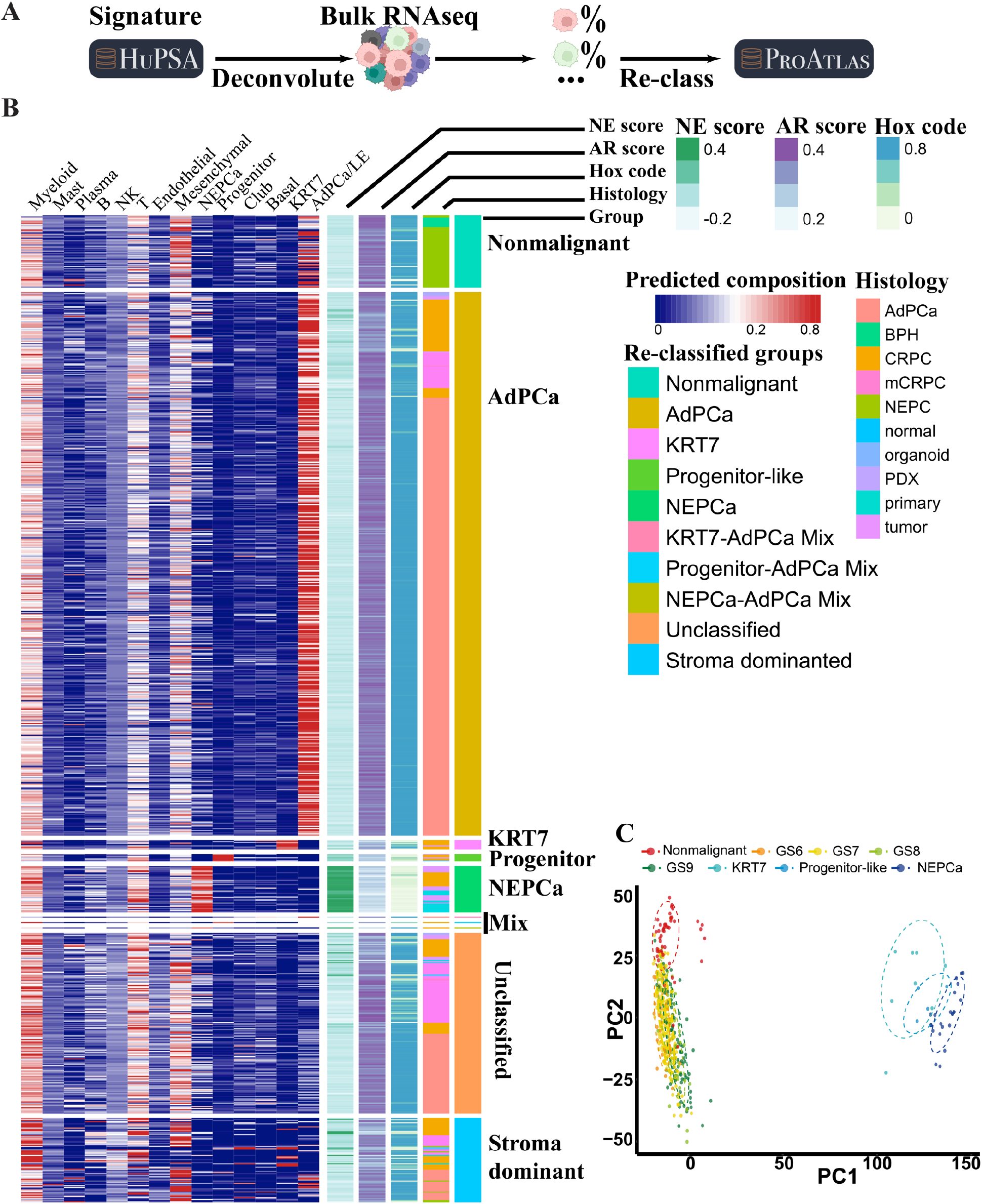
Establishment of ProAtlas through HuPSA-based deconvolution. (A) Workflow illustrating the process of establishing ProAtlas using HuPSA-based deconvolution. (B) Heatmap displaying the deconvolution result. Each PCa sample (n=877) was re-classified based on its molecular features defined by the HuPSA dataset. Samples were also annotated with NE score, AR score, correlation score with (prostatic) HOX code, and histology. (C) Principal component analysis (PCA) conducted on the ProAtlas samples. AdPCa samples were further classified based on their Gleason score information. The dashed circles represent the 75% data ellipse of the corresponding group.

Using the transcriptome profiles of individual populations in HuPSA, we conducted covariance-based deconvolution analysis on the ProAtlas collection. This analysis assessed the composition of various cell populations within the samples, including luminal, KRT7, basal, club, progenitor-like, NEPCa, stromal (fibroblast, smooth muscle, pericyte), and immune cells. The molecular subtypes were assigned to each sample based on its dominant populational composition.

As shown in Fig. 3B, most specimens can be re-classified into distinct subtypes, including luminal (n=570), KRT7 (n=9), progenitor-like (n=7), and NEPCa (n=43). The luminal group comprised both nonmalignant prostate and AdPCa tumors. Mixed groups and unclassified samples were also identified. Samples dominated by stromal cells contained minimal cancer cells. The reclassified samples were collected to construct the ProAtlas dataset.

Principal component analysis (PCA) of ProAtlas revealed clustering of samples of the same subtypes, indicating similar transcriptome profiles within each subtype (Fig. 3C). Notably, KRT7, progenitor-like, and NEPCa samples were distinctly separated from nonmalignant and AdPCa samples of various Gleason Scores, highlighting significant transcriptome differences in these aggressive PCa. This underscores the effectiveness of HuPSA deconvolution-based reclassification in accurately classifying PCa samples.

The analysis of ProAtlas data revealed distinctive features of KRT7 and progenitor-like PCa subtypes, characterized by low NE score, diminished AR score, and reduced prostatic HOX code (Fig. 3B). These findings suggest that both KRT7 and progenitor-like PCa subtypes represent de-differentiated double-negative prostate cancer, signifying a departure from the typical cellular characteristics associated with prostate adenocarcinoma.

Additionally, deconvolution analysis led to the identification of rare KRT7 and progenitor-like PCa samples in public study cohorts (Fig. 4A). Through collaboration, corresponding archived tissues from the GSE126078 cohort were obtained. All retrieved samples were initially diagnosed as DNPCa^31^. IHC staining was conducted to evaluate the expression of KRT7, FOXA2 and SOX2, the corresponding markers of the KRT7 and progenitor-like subtypes.

**Fig. 4.**
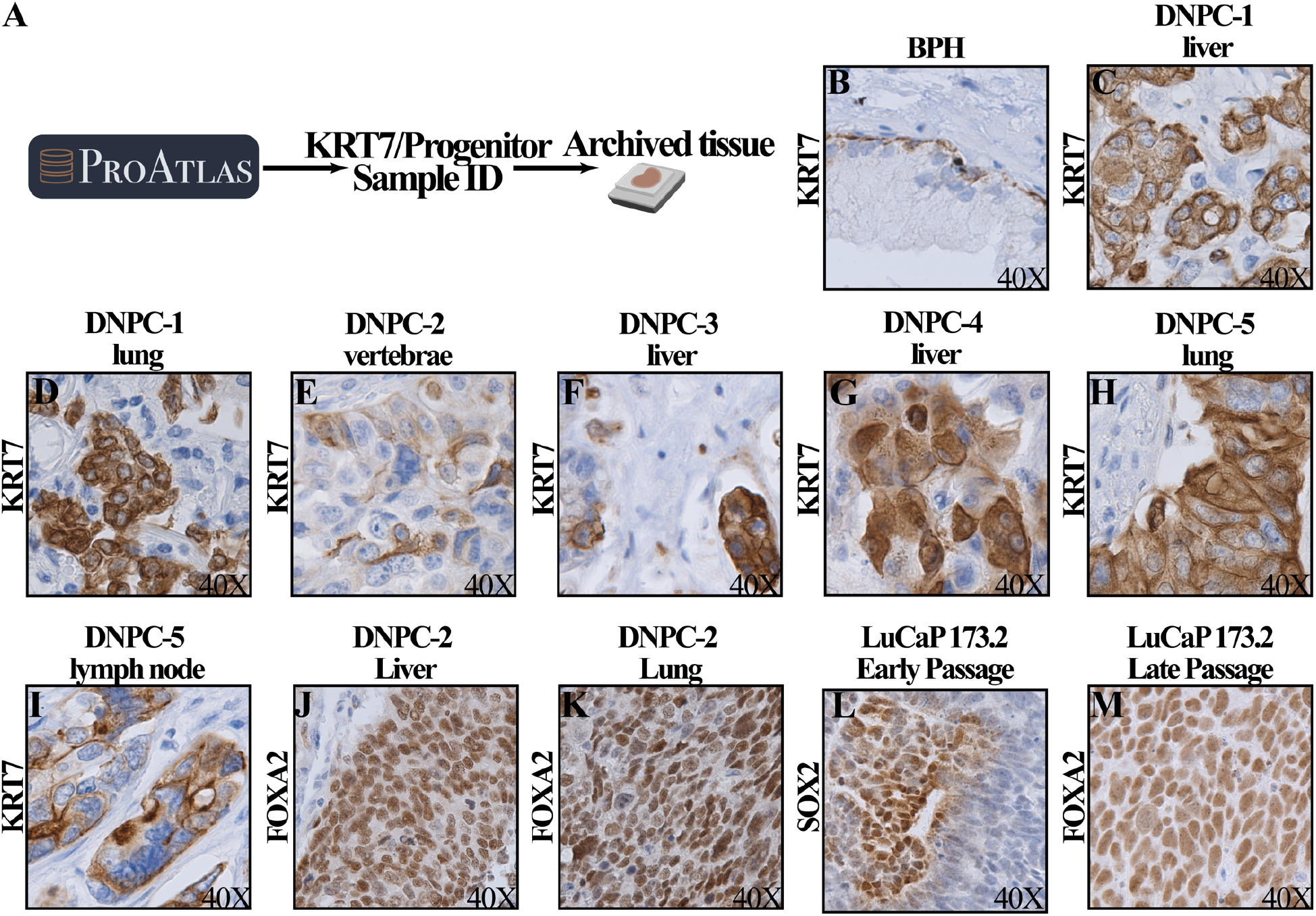
Validation of the novel KRT7 and progenitor-like PCa subtypes through immunohistochemistry staining. (A) The workflow chart illustrating the process of identifying KRT7 and progenitor-like patient samples in public cohorts. (B-I) Immunohistochemistry analysis demonstrated that KRT7 expression was exclusively detected in basal epithelial cells of benign prostatic hyperplasia (BPH) samples, whereas all deconvolution-identified samples of the KRT7 PCa subtype exhibited positive KRT7 protein staining. (J-M) Progenitor-like samples exhibited positive FOXA2 and SOX2 staining.

Remarkably, all seven DNPCa samples examined exhibited positive KRT7 staining in cancer cells (Figs. 4C-I). In contrast, KRT7 expression was absent in the luminal epithelia of benign prostatic tissues (Fig. 4B) and low-grade AdPCa specimens (data not shown, n>40).

Additionally, LuCaP-173.2 patient-derived DNPCa xenografts and three DNPCa metastases from the GSE126078 cohort were subjected to IHC evaluation of SOX2 and FOXA2 expression. Two of the three DNPCa metastases examined were positive for FOXA2 (Figs. 4J & 4K), and the expression of SOX2 and FOXA2 was detected in LuCaP 173.2 early-passage and late-passage tumors, respectively (Figs. 4L & 4M).

Notably, from DNPCa-2 patient, three samples were obtained, including vertebral, liver and lung metastases. From HuPSA-based deconvolution, the vertebral metastasis was reclassified as the KRT7 subtype, while the liver and lung metastases were reclassified as the progenitor-like subtype. This classification was validated by the positive IHC staining of subtype markers KRT7 and FOXA2 (Figs. 4E, 4J & 4K), providing additional validation for the accuracy of HuPSA deconvolution-based reclassification.

Collectively, the positive expression of KRT7, FOXA2, and SOX2 in DNPCa specimens serves as compelling evidence for the existence of these two novel subtypes in advanced PCa. This validation underscores the accuracy of the clustering approach employed in the construction of the HuPSA dataset, affirming its utility in delineating distinct PCa subtypes based on their molecular profiles.

### 5. Establishment of a public available web application for the visualization of gene expression in different subtypes of PCa

We previously introduced the PCTA web app, allowing researchers to visualize gene expression in commonly used cancer cell lines (https://pcatools.shinyapps.io/PCTA_app/)^32^. The positive reception highlighted a demand for such resources. In response, we developed the HuPSA-MoPSA web app, accessible at https://pcatools.shinyapps.io/HuPSA-MoPSA/. This platform enables users to visualize gene expression at single cell level by inputting official gene symbols. The app provides feature plots, violin plots and box plots, allowing simultaneous access to data from all three datasets established in this study, HuPSA, MoPSA and ProAtlas.

As illustrated in Fig. 5, users can input gene symbols to explore gene expression patterns across different datasets. High-resolution feature plots and violin plots are generated for HuPSA and MoPSA, while box plots are generated for ProAtlas, facilitating enhanced data interpretation and quantification.

**Fig. 5.**
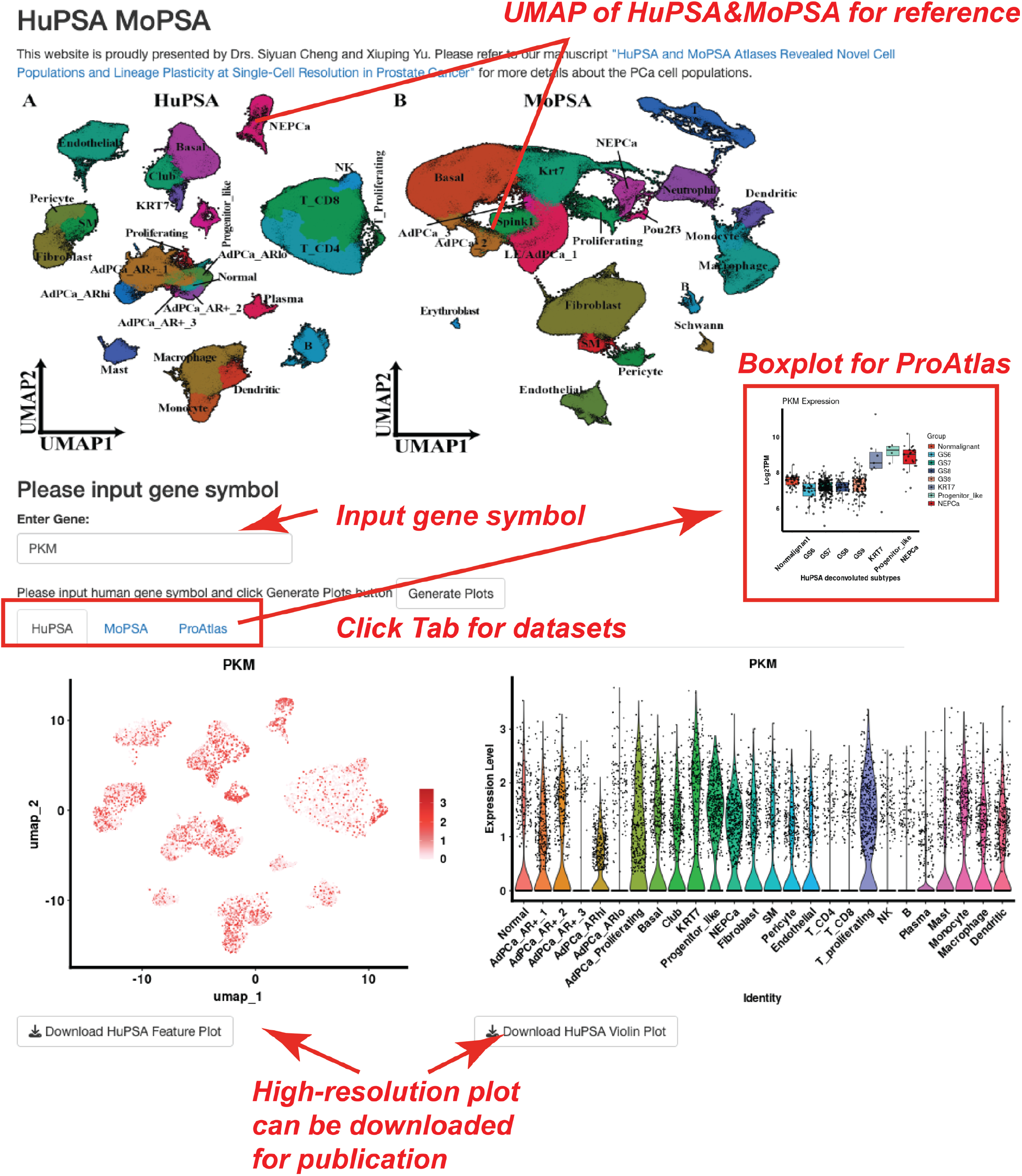
Demonstration of web application “HuPSA&MoPSA”. This app was developed for users to visualize and download data from HuPSA, MoPSA, and ProAtlas datasets established in this study. It allows users to explore genes of interest by inputting gene symbols. Human and mouse gene symbols are automatically converted for compatibility with different datasets. Under specific dataset tab, users can generate high-resolution, publication-level feature plots, violin plots, and box plots for detailed data analysis.

## Discussion

Advanced technologies have revolutionized our ability to understand pathological processes. RNA sequencing, for instance, provides unbiased and high-throughput gene expression profiling data. Single cell RNA sequencing technology enables researchers to investigate transcriptome at single-cell resolution, addressing old questions in cancer research and tackling new challenges such as phenotypic plasticity and non-mutational epigenetic reprogramming^33^. The integrated scRNAseq datasets, HuPSA and MoPSA, allows for the detection of various cell populations and visualization of cancer cells of different phenotypes on dimensional reduction maps. By analyzing differentially expressed genes and enriched signaling pathways, these atlases offer insights into the driver mechanisms of phenotypic plasticity and identify potential druggable targets.

In this study, we identified two novel DNPCa cell populations, progenitor-like and the KRT7 populations. Leveraging the compilation of multiple scRNAseq datasets, the establishment of HuPSA enabled the identification of these rare populations.

Progenitor-like cells, reminiscent of cancer stem cells, may contribute to tumor recurrence and therapy resistance. Characterized by the expression of SOX2 and FOXA2, genes also expressed in embryonic prostate and NEPCa cells, these cells represent a distinct PCa subtype. Our analysis revealed signature genes and enriched signaling pathways in this population, providing potential markers for defining this subtype. Previous studies have identified SOX2^+^ and FOXA2^+^ non-NEPCa cases through IHC staining, which likely represent the progenitor-like subtype of PCa^28,29^.

The KRT7 population, although located closely with basal epithelial cells on the UMAPs in both HuPSA and MoPSA, exhibits distinct gene expression patterns. Despite resembling basal cells on dimensional reduction maps, they lack canonical markers such as KRT5, KRT14, or TP63, instead highly expressing KRT7. This underscores the importance of exploring non-traditional markers in PCa classification.

Indeed, the expression of KRT7 (Cytokeratin 7), a type of intermediate filament protein, is typically found in glandular and transitional epithelial cells such as those in cervical epithelia. While KRT7 expression has been detected in various types of cancer, its presence in PCa cells has not been observed until now. Elevated KRT7 expression has been associated with aggressive cancer phenotypes including increased cell proliferation, migration, invasion, and epithelial-to-mesenchymal transition. ^34-36^ Moreover, in pancreatic cancer and gastric cancers, heightened KRT7 expression correlates with poor prognoses ^37,38^. The expression of KRT7 in double negative PCa cells implies a potential role for this protein in PCa progression.

Furthermore, utilizing HuPSA-based deconvolution analysis against bulk RNAseq data allowed for the re-classification of PCa samples. This approach identified rare samples of the newly defined progenitor-like and KRT7 subtypes, providing valuable insights into PCa heterogeneity.

Despite the advantages of integrated scRNAseq datasets, such as HuPSA and MoPSA, they come with limitations. The complexity of scRNAseq data, represented by multi-dimensional gene expression profiles, poses challenges in data visualization and interpretation. Dimensional reduction techniques may not fully capture transcriptome identities, and interpretation can be subjective. As a famous ancient Chinese poet Su Shi eloquently put it, “It’s a range viewed in face and peaks viewed from the side, assuming different shapes viewed from far and wide.” Objective data processing is crucial, but researchers should remain aware of potential biases in data interpretation.

## Methods

### 1. Single-cell RNA sequencing on TRAMP tumors

TRAMP tumor tissues were harvested and dissociated using “Tumor Dissociation Kit, mouse, #130-096-730” from Miltenyi Biotec. The dead cells were removed using “Dead Cell Removal Kit, #130-090-101” from Miltenyi Biotec. The red blood cells were removed using “Red Blood Cell Lysis Solution, # 130-094-183” from Miltenyi Biotec. The cDNA libraries were constructed using Chromium Next GEM Single Cell 3’ Reagent Kit v3.1. The resulting cDNA was fragmented, processed, and size selected. Adaptors were ligated and dual indexes were attached via PCR. The libraries were cleaned up and assessed for quality with the Agilent TapeStation D5000 High Sensitivity Kit. The libraries were quantitated with qPCR (NEBNext Library Quant Kit for Illumina, Bio-rad CFX96 Touch Real-Time PCR) and normalized to 1.5 pM and pooled. The library pool was denatured and diluted to approximately 300 pM. A library of 1.5 pM PhiX was spiked in as an internal control. Paired-end sequencing was performed on an Illumina NovaSeq 6000.

### 2. Public scRNAseq collection and alignment

The human and mouse scRNAseq data were collected from Gene Expression Omnibus (GEO). Only 10X genomic based single-cell studies were considered. Raw data of 6 human studies in the SRA database were used to build HuPSA, including GSE210358^6^, GSE137829^13^, GSE193337^14^, GSE206962, GSE181294^15^, GSE185344^16^ . Data of 12 mouse studies were used to build MoPSA, including GSE146811^39^, GSE164969^17^, GSE201956^40^, GSE216158^18^, GSE221755^19^, GSE171336^20^, GSE165741^21^, GSE174471^22^, GSE210358^6^, GSE163316^23^, GSE189307^24^, GSE158468^25^, GSE153701^26^ (sTable4). All FASTQ files were downloaded using SRA-Toolkit fasterq-dump function and aligned to human (GRCh38 from NCBI) or mouse (GRCm38 from Ensembl) genome using Alevin-fry^41^. The nf-core/scrnaseq pipeline was used for software environment, paralleling and quantity control. Linux ubuntu operating system was used for data analysis.

### 3. scRNAseq Data integration and processing

Single-cell read count data were loaded into R environment using “loadFry” function from Fishpond package^42^. “Seurat V5”^43^ was used for quality control, data integration, UMAP dimensional reduction, clustering and annotation. Quality control was performed based on the total RNA count and feature count for each cell. Cells with extremely low or high RNA/feature counts were excluded to ensure the removal of low-quality cells and doublets. The RPCA model was used for data integration. In the integration process, each study (instead of sample) was treated as one “batch” to prevent over integration. After the UMAP dimensional reduction and clustering, the clusters were then labeled with population names based on the marker genes expression and original histology information. The annotated human lung V2 (HLCA) dataset from “Azimuth”^44^ were used as reference for the manual annotation of immune and stromal populations. The R package “SCpubr”^45^ was used for data visualization. Dim plots were utilized to visualize the location on UMAP of each cell population in HuPSA and MoPSA. Feature plots were employed to visualize the gene expression on UMAPs. Geyser plots were used to compare gene expressions among the different cell populations. In Geyser plot, the dots along the y-axis represent the distribution of gene expression levels in single-cells. The black balls indicate the median values of each cell population. The ends of thick line and thin line indicate 66% and 95% of the data points within each group, respectively. Sankey plots were used to depict the data “flow” among populations, histology, and study groups. The single cell copy number alterations were calculated and visualized using “InferCNV” package^46^. Pathway activity was calculated using “decoupleR”^47^. Transcription factor activity was calculated using “dorothea”^47^. The “CSCDRNA” package^30^ was used for HuPSA-based bulk RNAseq deconvolution. The “CellChat” package^48^ was used to predict ligand-receptor pairs in HuPSA and MoPSA. All detailed parameters were described in the R scripts deposited to GitHub.

### 4. Bulk RNAseq data collection and processing

Bulk RNAseq samples were selected based on the following criteria: duplicated reads < 60%, total reads > 20 million, and STAR unique mapping > 70%. However, the TCGA samples were not filtered because these specific metrics were not available. Eventually 877 high quality samples comprising normal, BPH, tumor, primary, AdPCa, CRPCa, mCRPCa, NEPCa, organoid, PDX (patient-derived xenograft) histology groups were selected and used for HuPSA-based deconvolution analysis. The FASTQ files were collected from multiple cohorts including GSE120795^49^, GSE126078^31^, GSE156289^50^, GSE114740, GSE217260, GSE179321^51^, GSE112786^52^, GSE104131^53^, GSE31528^54^, GSE54460^55^, GSE22260^56^, GSE216490^57^; dbGaP accessions: phs000909.v1.p1^58^, phs000915.v2.p2^2^. The TCGA_PRAD read counts and TPM data were acquired using “recount3”^59^ R package. The sequencing reads were aligned to human GRCh38 genome (from NCBI) using STAR^60^ and Salmon^61^. The nf-core/rnaseq pipeline was used for paralleling and quantity control.

### 5. Human prostate specimens

De-identified human prostate tissues (archived) were obtained from LSU Health-Shreveport Biorepository Core, Overton Brooks VA Medical Center, and Genitourinary Cancer Specimen Biorepository at University of Washington. All the tissues were used in accordance with the protocols approved by the Institutional Review Board of LSU Health-Shreveport.

### 6. Immunohistochemical, immunofluorescence staining and Western blot

Primary antibodies used in immunostaining include KRT7 (abclonal, A4357), MMP7 (abclonal, A0695), CK5 (BioLegend, 905904), CK8 (BioLegend, 904801), SOX2 (Cell signaling, 3728), FOXA2 (Abcam, 108422). IHC staining was performed using Vectastain elite ABC peroxidase kit (Vector Laboratories, Burlingame, CA). The tissue sections were counterstained, mounted, and imaged with a Zeiss microscope (White Plains, NY).

For Western blot, cells were collected in PBS and lysed in passive lysis buffer (Promega, Madison, WI). Equal amounts of protein were loaded for Western blot analyses. ProSignal Dura ECL Reagent (Genesee Scientific, San Diego, CA) and Chemidoc (Bio-Rad, Hercules, CA) were used to visualize the protein bands.

## Acknowledgement

RNA sequencing data were downloaded from dbGaP through dbGaP accession number phs000909 and phs000915^2,62^. The data were generated under the support of NHGRI grant # U54 HG003067 to E. Lander at Broad Institute and SU2C/PCF Prostate Dream Team Translational Cancer Research Grant.

## Declarations

### Authors’ contributions

Conceptualization, Investigation, Methodology: SC, XY; Data curation: SC, LL; Software, Visualization, Writing – original draft: SC; Formal analysis: YY, YS; Resources, Funding acquisition, Supervision: XY; Writing – review & editing: QG, OF, EC, XY.

---

**sFig 1.**
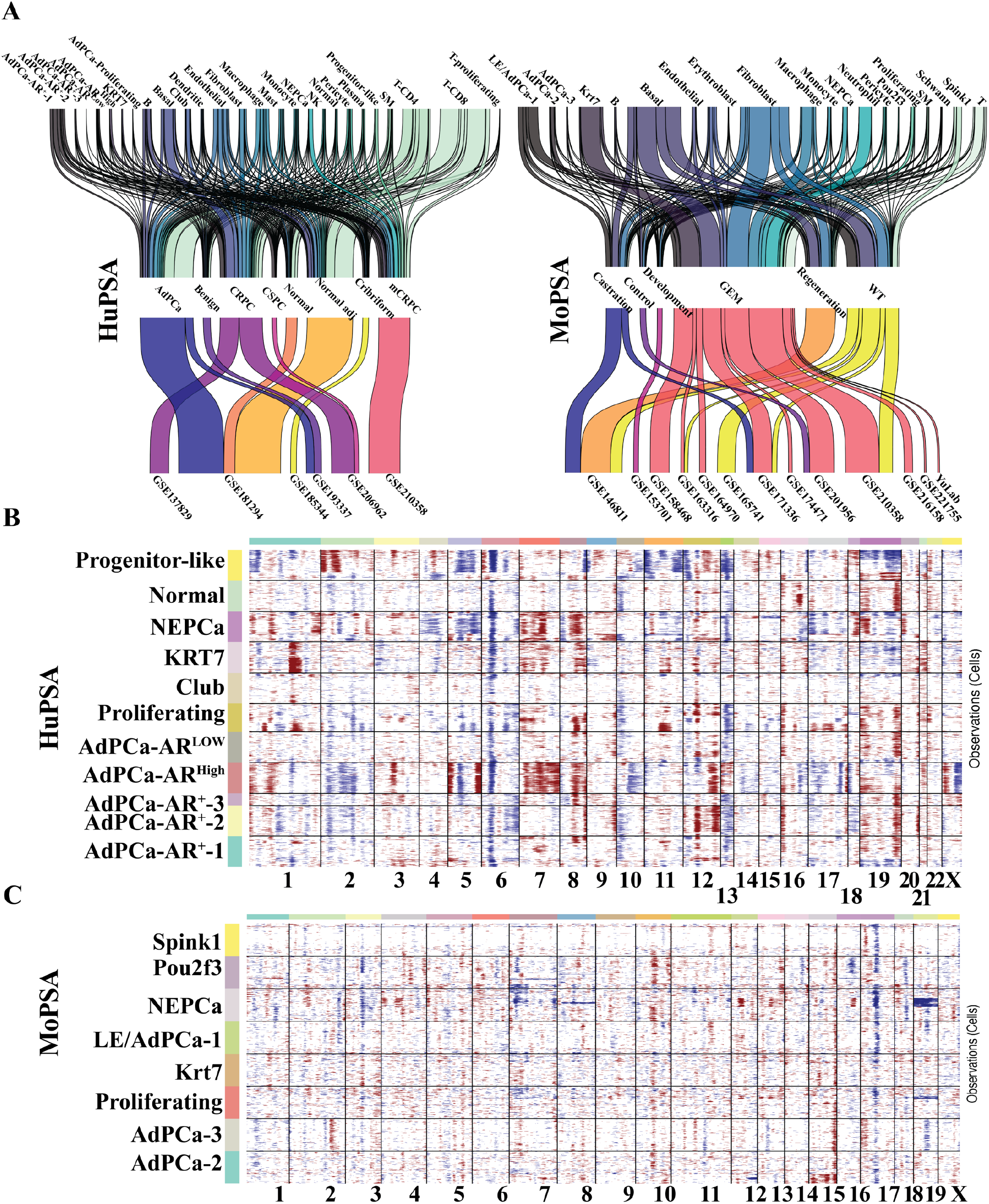
Sankey plots of HuPSA (A) and MoPSA (B) to depict the data flow from the newly-defined cell populations (left) to histology groups (middle) and the studies that collected these data (right). (C) InferCNV analysis of the epithelial populations in both HuPSA (up) and MoPSA (down).

**sFig 2.**
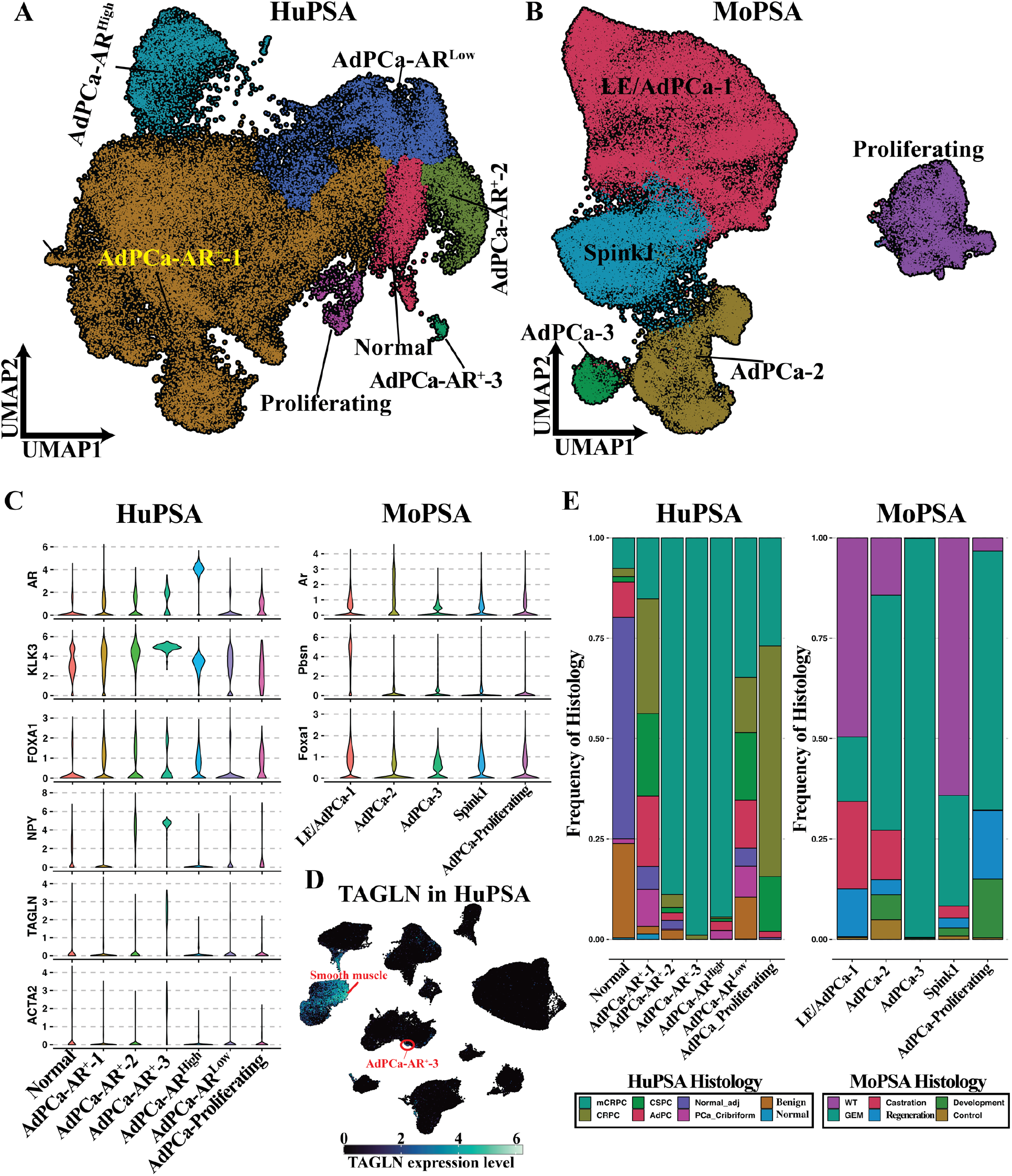
UMAPs of Luminal/AdPCa populations in HuPSA (A) and MoPSA (B). (C) Violin plots to visualize genes expression in the Luminal/AdPCa populations in HuPSA (left) and MoPSA (right). (D) Feature plot to visualize the expression of the TAGLN gene in HuPSA. TAGLN is expressed in both smooth cells and AdPCa-AR+-3 cells. (E) Bar plots to depict the sample composition of each Luminal/AdPCa populations in HuPSA (left) and MoPSA (right).

**sFig 3.**
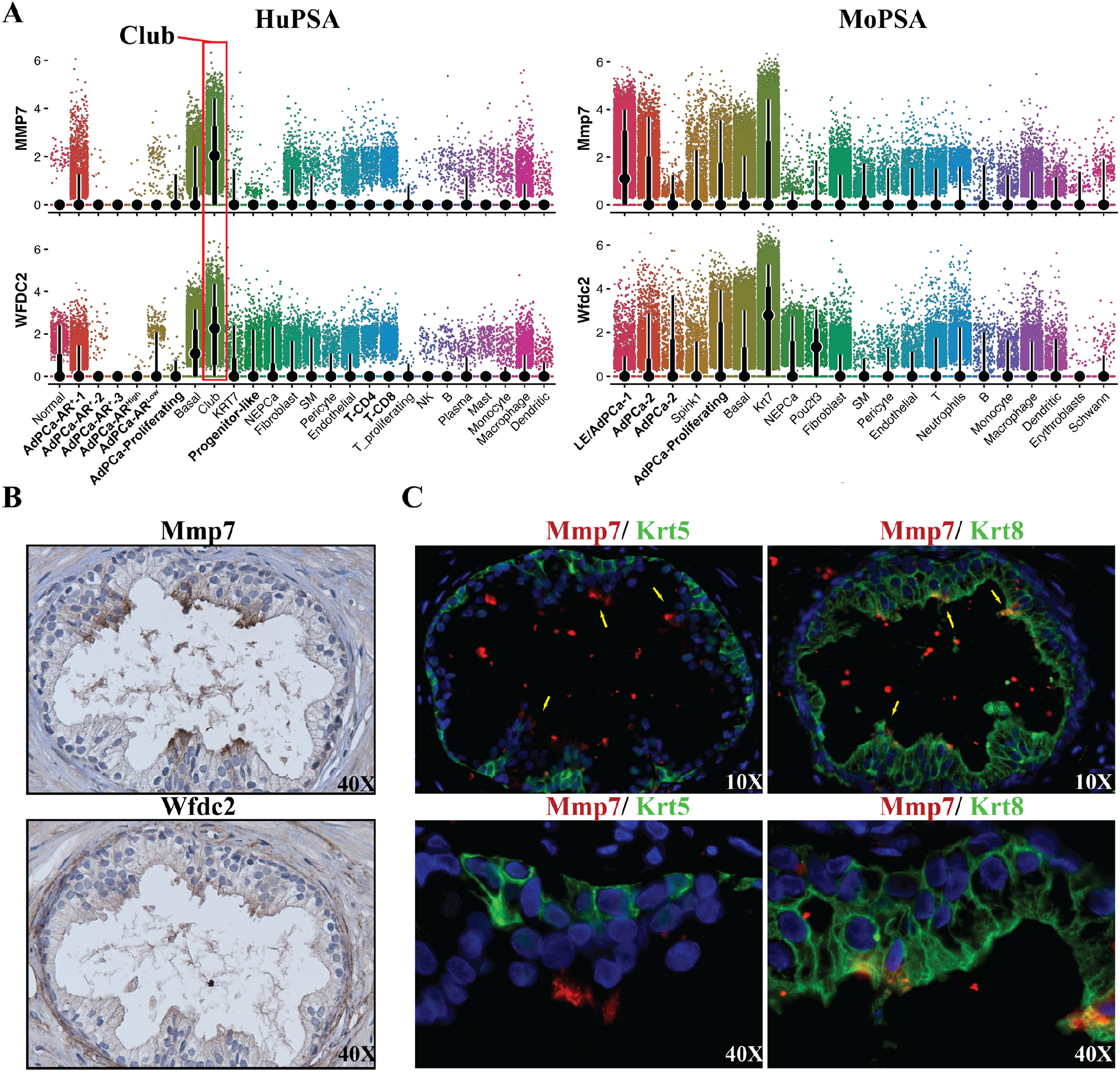
Club population. (A) Geyser plots to demonstrate the co-expression of MMP7 and WFDC2 in the Club cells of HuPSA. (B) Immunohistochemistry staining of Mmp7 and Wfdc2 on serial sections confirms the presence of Club cells in human BPH samples. (C) Dual immunofluorescence staining of Mmp7 with luminal epithelial marker Krt8 or basal epithelial marker Krt5 indicates that Mmp7+ cells are luminal but not basal epithelia.

**sFig 4.**
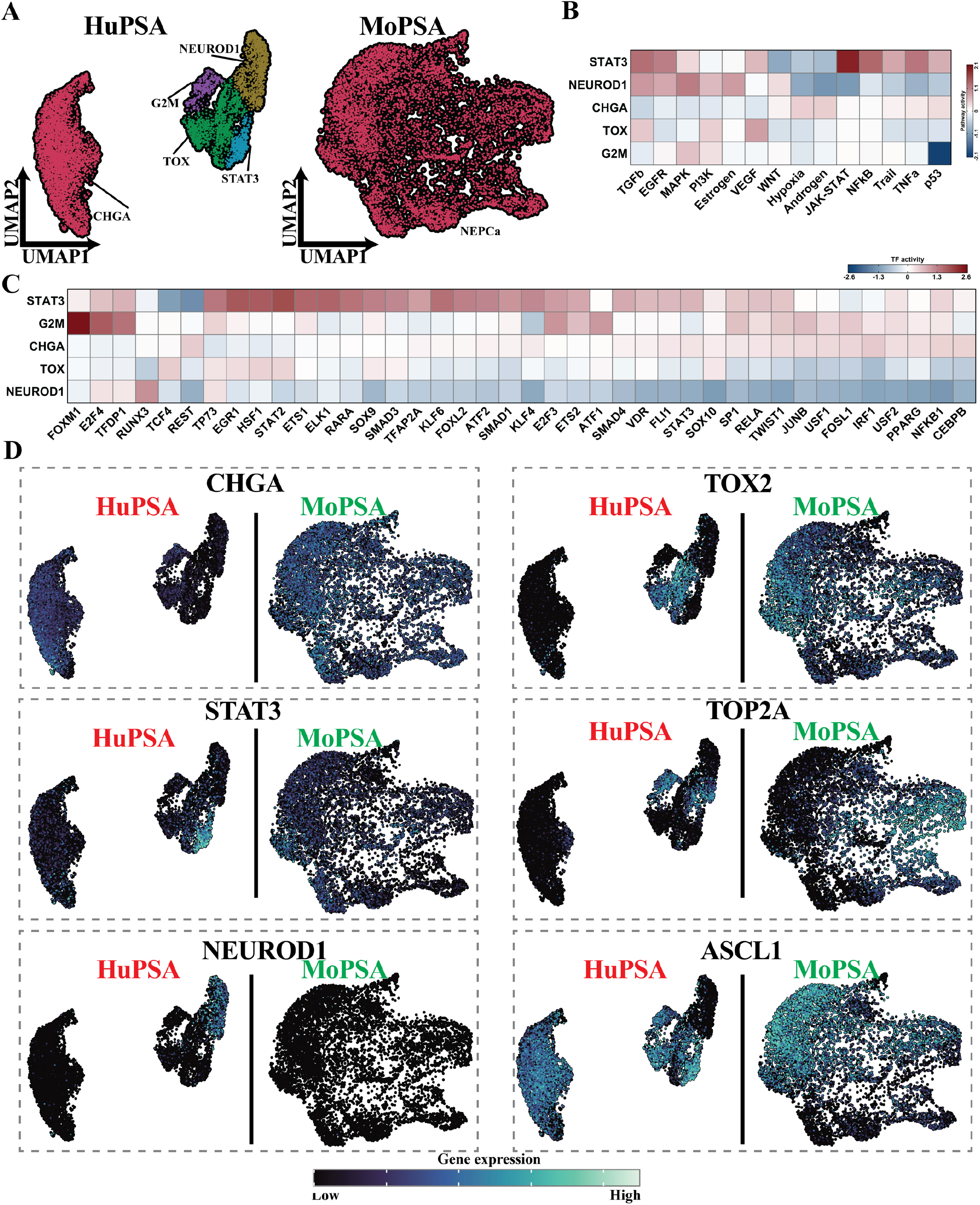
NEPCa populations. (A) The UMAP plots revealed heterogenous NEPCa cells in HuPSA and homogenous NEPCa cells in MoPSA. Five NEPCa sub-populations were identified in HuPSA including CHGA, TOX, G2M, STAT3 and NEUROD1. (B & C) Heatmaps illustrate the activities of enriched signaling pathways and upstream transcription factors for the NEPCa sub-populations in HuPSA. (D) Feature plots visualize the expression of marker genes in the NEPCa cells of both HuPSA and MoPSA datasets. It is noteworthy that the expression of ASCL1 and NEUROD1 is exclusive in human NEPCa cells.

