## Supplementary material for "Unveiling Novel Double-Negative Prostate Cancer Subtypes Through Single-Cell RNA Sequencing Analysis": Supplimental information

Supplemental information

1. Charactering the previously known populations within the HuPSA and MoPSA datasets

The luminal populations were defined by the expression of prostatic luminal epithelial marker genes FOXA1, HOXB13, and NKX3-1 (Fig. 1C). The transcriptomes of luminal cells were extracted from HuPSA and MoPSA datasets and visualized on UMAPs for higher resolution (sFig. 2A & 2B). In HuPSA, the luminal cells consisted of seven populations including normal (contributed by normal, normal-adjacent, benign, and some AdPCa samples), three AdPCa-AR^+^ groups, AdPCa-AR^high^, AdPCa-AR^low^, as well as highly proliferating groups. The AdPCa-AR^+^-1 population only slightly deviated from normal cells on UMAP (sFig. 2A). Its inferred copy number variation (iCNV) pattern was also similar to that of normal cells (sFig. 1B). This population may represent low grade PCa, and cells in this population are still at the early stage of transformation. In contrast, the AdPCa-AR^+^-2 population harbored higher level of iCNVs (sFig. 1B). The highest iCNVs level was present in the AdPCa-AR^high^ population (sFig. 1B), suggesting a genome instability in these cells. The AdPCa-AR^+^-2 and AdPCa-AR^high^ populations were mostly derived from advanced PCa samples including CRPCa, mCRPCa, and CSPCa (sFig. 2E). Another population that was mostly derived from advanced PCa was AdPCa-AR^+^-3 (sFig. 2E). It departed from the main AdPCa cluster (sFig. 2A), but its iCNV pattern did not significantly differ from that of normal cells (sFig. 1B), suggesting that genome instability may not be a driving mechanism of cancer progression in these cells. It is interesting to note that the AdPCa-AR^+^-3 population expressed NPY, a gene widely expressed in the central nervous system, as well as two smooth muscle markers TAGLN and ACTA2 (sFigs. 2C & 2D). This suggests that while still expressing luminal genes such as HOXB13, FOXA1 and NKX3-1, AR^+^-3 population may have acquired certain mesenchymal features. These cancer cells are likely driven by epigenetic reprogramming.

The AdPCa-AR^low^ population, although low in AR expression, still expressed KLK3 and NKX3-1, two AR target genes, at fair levels (sFig. 2C), indicating an active AR signaling in these cells.

The proliferating population exhibited high proliferative activity as indicated by the high percentage of cells at predicted G2M/S phases of the cell cycle as well as the high expression of MKI67 and TOP2A (Fig. 1C). The cells in this population were mostly derived from CRPCa and mCRPCa as well as CSPCa samples (sFig. 2E). This population exhibited active AR signaling, indicated by the expression of AR and AR target KLK3 (aFig. 2C).

The luminal populations in MoPSA include­­d LE/AdPCa -1, AdPCa -2 & -3, and highly proliferating populations, as well as a unique Spink1 population that was not detected in HuPSA (sFig. 2B). The Spink1 population was predominantly contributed by the prostates of wild-type mice (sFig. 2E) and it harbored low iCNVs (sFig. 1C). Because of its absence in HuPSA and the relatively normal molecular profiles, the Spink1 population will not be further characterized in this study. All the other luminal cells in MoPSA expressed Hoxb13 and Foxa1 (Fig. 1C), like those in HuPSA. But NKX3-1, a prostate differentiation marker, was only expressed at high levels in the normal LE/AdPCa-1 population (Fig. 1C), suggesting a loss of differentiation state in the other populations. In addition, unlike the luminal populations in HuPSA, all of which expressed the AR target genes (such as KLK3) regardless of AR expression status, only LE/AdPCa-1 but not any other luminal populations in MoPSA expressed AR target gene, Pbsn (sFig. 2C). Given that the LE/AdPCa-1 population predominantly originated from wild-type mice, with only a limited number of cells originating from genetically engineered mice, and it harbored low iCNVs (aFig. 2E & sFig. 1C), this population may represent normal prostate luminal epithelia (LE) and the cells at an early stage of transformation. The other AdPCa populations, including AdPCa -2 & -3 and highly proliferating populations, on the other hand, were predominantly contributed by GEM (sFig. 2E) and harbored slightly higher level of iCNVs than the LE/AdPCa-1 population (sFig. 1C).

2. MMP7 and WFDC2 are novel markers for prostate club cells

The prostate club cell population was observed in HuPSA but not in MoPSA. This population, recently identified in the prostate through scRNAseq analysis, was named after airway club cells in the lung, a dominant secretory cell population characterized by the expression of the SCGB1A1 and MMP7^27,36^. In the prostates, similar to the KRT7 population, club cells were clustered together with basal cells on UMAP but did not express basal epithelial markers (sFig. 3A & Fig. 1C). Instead, they expressed markers of airway club cells including MMP7, WFDC2 and SCGB1A1 (sFig. 3A).

The HuPSA club population exhibited low iCNVs (sFig. 1C), suggesting that these cells are likely non-cancer cells, consistent with literatures^36^. As illustrated in sFig. 3A, MMP7 was expressed in the club population, whereas WFDC2 was expressed in both club and basal populations. This expression pattern was confirmed by IHC staining. MMP7 expression was detected in in the apical side of a subset of luminal epithelial cells in human benign prostatic hyperplasia (BPH) samples (sFig. 3B). On the serial section, WFDC2 expression was detected in the basal cells as well as in the areas positive for MMP7 immunostaining (sFig. 3B). Moreover, dual immunofluorescence staining results showed that MMP7 was co-expressed with luminal marker KRT8 but not basal marker KRT5 (sFig. 3C), indicating that the club population, although clustered together with basal cells, are luminal epithelial cells.

3. Mouse NEPCa uniformity vs. human NEPCa subpopulations

NEPCa cells from both HuPSA and MoPSA expressed NEPCa markers INSM1 and NCAM1 (Fig. 1C). For a more detailed analysis of NEPCa cells, the transcriptomes of these cells were extracted from the HuPSA and MoPSA datasets, and subsequently visualized on UMAPs (sFig. 4A). While the mouse NEPCa population was relatively uniform, the human NEPCa cells can be further divided into distinct sub-populations. Through unsupervised clustering and examination of marker genes’ expression, the HuPSA NEPCa cells could be further divided into five sub-populations based on the expression of marker genes or the pathways enriched, including CHGA, G2M, TOX, STAT3, and NEUROD1 (sFigs. 4A-4C). It is noteworthy that the expression of marker genes was not mutually exclusive among these groups.

The CHGA subpopulation expressed high levels of the NE marker CHGA (sFig. 4D). It represents the canonical NEPCa phenotype. The other NEPCa sub-populations expressed relatively low but non-negligible levels of CHGA.

The G2M sub-population was characterized by the high expression of the signature genes associated with cell cycle G2 & M phases such as TOP2A, MKI67, CENPF/W, CKS1B, CKS2 and STMN1. Concurrently, this sub-population displayed heightened transcriptional activity of FOXM1 (Fig. 6C), a transcription factor linked to cell proliferation. In contrast, the other subpopulations exhibited an even distribution of genes related to G1, S and G2M phases. Notably, multiple genes enriched in this sub-population are involved in DNA damage repair (DDR) pathway, including ASPM, CDKN2A/D, CDKN3, CKAP2, ECT2, NUCKS1, PTTG1, HMGB1-3, HMGN2, and TPX2, suggesting an increased DDR activity. Additionally, the G2M subpopulation was predicted to maintain a low P53 signaling activity (sFig. 4B), highlighting its distinct properties compared to other NEPCa subpopulations.

The TOX sub-population was distinguished by the high expression of Thymocyte selection-associated HMG box gene (TOX). These cells exhibited heightened activity in the VEGF signaling pathway and expressed elevated levels of TCF4, a transcription factor of Wnt signaling (sFigs. 4B-D).

The STAT3 sub-population not only expressed elevated levels of STAT3 mRNA but also displayed heightened activity in the JAK-STAT signaling, along with increased transactivation of STAT3 target genes (sFigs. 4B-D). Active JAK/STAT signaling is known to promote cellular plasticity^6^, suggesting that the STAT3 subpopulation could be driven by JAK-STAT3 signaling.

The NEUROD1 sub-population was characterized by the elevated expression of NEUROD1, GRID2 and GKAP1. It exhibited heightened activities in TGF, EGFR, MAPK, PI3K, ER, and Wnt signaling pathways (sFig. 4B). The enrichment of Wnt signaling in this subpopulation aligns with a previous study demonstrating that Wnt signaling activates NEUROD1 expression during neurogenesis in mice^37^. In contrast, the expression of ASCL1, a driver transcription factor of NEPCa^38^, was detected in all the NEPCa subpopulations except NEUROD1. By examining the feature plots of NEUROD1 and ASCL1 in HuPSA, we observed a mutually exclusive expression pattern between NEUROD1 and ASCL1 in NEPCa (sFig. 4D).
